# The Pulmonary Microbial Signatures of Early Stages of Severe Infection in Children

**DOI:** 10.1101/2025.08.25.672262

**Authors:** Hao Wu, Yinghu Chen, Dengming Lai, Wei Li, Mingming Zhou, Fang Shu, Chen Yuan, Ping Guo, Weijie Yu, Xiaofeng Chen, Xuan Nie, Ran Tao, Lingfeng Mao, Qiang Shu, Shiqiang Shang

## Abstract

The respiratory tract related diseases are the main causes of death in children. Child pulmonary severe infection is the most important inducement for Respiratory tract related diseases. Therefore, it is necessary to understand the composition of the microbial community and the species correlation in children with severe pulmonary infection. In this study, the bronchoalveolar lavage fluid (BALF), blood and cerebrospinal fluid (CSF) samples of 782 children with severe infection in the early stage were systematically analyzed by amplicon-sequencing technology to reveal the distribution of microbial community and its clinical correlation. Results reveal significantly higher bacterial abundance in BALF than in blood/cerebrospinal fluid, with community structures shaped by tissue microenvironments, geographical disparities, age, and clinical symptoms. Neonatal BALF harbors simplified microbiota dominated by probiotics, while diversity increases with age, showing marked differences between 1-3 and 6-12-year-olds. Sepsis samples exhibit reduced microbial diversity, with *Staphylococcus* and *Moraxella* enrichments in the CST-4 cluster. Respiratory symptom progression correlates with microbiota succession from oral colonizers to respiratory pathogens. Third-generation sequencing-derived co-occurrence networks illustrate synergistic interactions among opportunistic pathogens, providing a basis for ecological targeted therapy. This study provides a key basis for breaking through the limitations of traditional diagnosis, establishing a precise diagnosis and treatment system based on third-generation sequencing, and developing ecological targeted treatment strategies.

## Introduction

Pneumonia or lower respiratory tract infection causes considerable morbidity and mortality in children worldwide, accounting for nearly 12% of deaths in Pediatric Intensive Care Units (PICUs)[1, 2]. The epidemiological characteristics of severe lung infection children (especially neonatal, infant and children with immune deficiency) because of immature immune system, the respiratory tract anatomy physiology characteristic (such as airway stenosis, cilia clearance ability), is a lung infection of high-risk groups. Severe pulmonary infection (such as severe pneumonia, lung abscess, sepsis-associated lung injury, etc.) has an acute onset and rapid progress, which is easy to cause respiratory failure, multiple organ dysfunction, and even death. Common pathogens include viruses (such as respiratory syncytial virus, influenza virus, adenovirus), bacteria (such as Streptococcus pneumoniae, Staphylococcus aureus), fungi (such as Candida, Aspergillus) and atypical pathogens (such as Mycoplasma pneumoniae, Chlamydia), and the proportion of mixed infections is high, which increases the difficulty of diagnosis[3]. The early diagnosis of clinical demand urgency children serious lung infection is closely related to the prognosis and treatment time. Delayed or inappropriate antimicrobial therapy, such as blind use of broad-spectrum antibiotics, may lead to the growth of drug-resistant bacteria, dysbiosis of flora, and even aggravation of the disease. Especially in children with immunosuppression (such as tumor chemotherapy and congenital immunodeficiency), the pathogens may be rare or opportunistic pathogens (such as cytomegalovirus and Pneumocystis carinii). Early identification of pathogens is the key to saving lives.

Traditional microbiological detection methods, such as sputum culture, blood culture, and pleural effusion culture, rely on pathogen culture, but children have low cooperation (it is difficult to obtain qualified sputum samples), long culture period (usually takes 48-72 hours), and the positive rate is affected by the use of antibiotics, sample contamination, and other factors, which may easily lead to treatment delay. In addition, although imaging, such as X-ray and CT, can show lung lesions, it cannot determine the type of pathogen, and it is difficult to guide accurate treatment. In recent years, the development of molecular biology techniques, such as multiplex PCR, metagenomic next-generation sequencing mNGS, third-generation target sequencing tNGS, pathogen specific antigen/antibody detection, provides the possibility for early and rapid detection. These technologies can break through the limitations of traditional culture and directly detect pathogen nucleic acid or antigen to achieve "precise pathogen diagnosis", especially for point-of-care rapid detection of severe infections. Early identification of pathogens, such as distinguishing bacterial and viral infections, can avoid unnecessary use of antibiotics and reduce the risk of drug resistance, such as reducing the blind use of β-lactam antibiotics in Mycoplasma pneumoniae infection. At the same time, targeted therapy for pathogens (such as antiviral drugs and narrow-spectrum antibiotics) can accelerate the remission of the disease, reduce complications (such as atelectasis and empyema), and reduce medical costs. The mortality rate of severe pneumonia in children is still high in developing countries (about 5%-15%), and early microbiological diagnosis can avoid delays in treatment. For example, in children with invasive pulmonary aspergillosis, early antifungal treatment reduces mortality from more than 50% to 20%-30%; For children with influenza virus and bacterial infection, timely combination of antiviral and antibiotic therapy can significantly improve the survival rate. Early diagnosis can identify special pathogens and mixed infections. In children with hematopoietic stem cell transplantation and congenital immunodeficiency, pulmonary infection may be caused by rare pathogens such as *cytomegalovirus* and *Pneumocystis carinii*, which are difficult to detect by routine culture. At the same time, children’s pulmonary infections are often mixed with multiple pathogens (such as bacteria-fungi mixed infection). Early identification of the infection spectrum can guide the combination of drugs and avoid the limitations of single treatment.

The third-generation sequencing technology (such as PacBio single-molecule real-time sequencing and Oxford Nanopore sequencing) has unique value in the early diagnosis of severe infection in children due to the characteristics of ultra-long read length and Nanopore without PCR amplification, combined with targeted enrichment technology to form tNGS. Its full-length gene precision analysis ability can directly read the complete gene sequence of pathogens to avoid the splicing error of second-generation sequencing, realize the accurate identification of strains/subtypes (such as distinguishing Klebsiella pneumoniae ST11 and ST258 types), and can also sequence the full-length drug resistance genes to guide the selection of antibiotics. The long-read advantage can solve the common problem of mixed infection of homologous bacteria in children and directly distinguish different strains of the same species or different strains of the same species. Nanopore technology supports sequencing and analysis at the same time, with a preliminary report in as soon as 2-4 hours. It is suitable for emergencies such as septic shock that need to start precision treatment within 6 hours. In addition, it can enrich nucleic acid without PCR amplification for trace samples (such as 100μl alveolar lavage fluid, 200μl blood) in children to reduce missed detection.

Bacterial 16S rRNA full-length sequencing and fungal ITS sequencing based on nanopore third-generation sequencing technology can more deeply analyze the microbial characteristics of disease samples. Studies have analyzed the lower respiratory tract microbiota of children with mycoplasma pneumonia (MPP) by 16S rRNA gene sequencing and found that the abundance of *Daulinella* and *Mycoplasma* in MPP group was significantly increased (over 67% and 65%, respectively), and the αdiversity of microbiota in severe MPP group was lower and the abundance of Mycoplasma was higher. Its abundance was positively correlated with complications and clinical indicators, revealing the characteristics of the lower respiratory tract microbiota and its association with disease severity in MPP[4]. Another study analyzed the bacterial community of bronchoalveolar lavage fluid in 48 infants by 16S sequencing and found that the microbial diversity of the lower respiratory tract in the recurrent wheezing group (A1 and A2) was significantly lower than that in the control group (B). The abundance of Sphingomonas and *Phyllobacter* in the A1 group, and *Phyllobacter* in the A2 group was significantly higher than that in the B group. The abundance of Prevotella, Neisseria and Haemophilus in group B was higher than that in the wheezing group. The α/β diversity analysis based on 6644 OTU (39.25% genus level annotation) showed significant differences between groups. It is suggested that recurrent wheezing in infants and young children may be related to the changes of overall bacterial microecology of the lower respiratory tract and the destruction of host respiratory immune homeostasis [5]. In the point-of-care rapid detection and dynamic monitoring, the MinION portable sequenator can be used for real-time sequencing of alveolar lavage fluid in ICU bedside, achieving rapid diagnosis within 3 hours of "sample intake-result", and guiding adjustment of anti-tuberculosis and other treatment programs by changes in pathogen load and clearance of drug resistance genes.

Early microbiological diagnosis of severe pulmonary infection in children is the core link to break through the limitations of traditional empirical treatment and achieve precision medicine. The background is due to the high susceptibility of children, the complexity of pathogens and the lag of traditional diagnosis, while the importance is reflected in many aspects such as precision treatment, improved prognosis, drug resistance control and public health. Therefore, it is necessary to understand the composition of the microbial community and the species correlation in children with severe pulmonary infection. The aim of this study is to explore the characteristics of pathogenic bacteria, non-pathogenic bacteria and opportunistic bacteria in the early stage of severe pulmonary infection in children by analyzing the composition and structure of microbial communities in bronchoalveolar lavage fluid (BALF), blood and cerebrospinal fluid (CSF) of more than 500 children with severe pulmonary infection in different children’s hospitals in Hang-Jia-hu area by amplicons sequencing. To provide data support for early warning and early infection control of severe infection in children.

## Material & Method

### Study participants and sampling

All the patients involved in this study were children being newly diagnosed as the respiratory tract related diseases at mild, moderate and severe phases and admitted to the Department of Respiratory Disease, Children’s Hospital in Hangzhou, Jiaxing, Huzhou, Shaoxing, and Jinhua areas. The symptoms of respiratory tract related diseases were based on clinical presentations including cough, fever, bronchitis, bronchopneumonia, pneumonia, as well as sepsis. Bronchoscopy with BAL collection was performed in accordance with the European Respiratory Society guidelines [6], as described previously [7]. Briefly, the collection of bronchoalveolar lavage fluid for children should be carried out under sedation or anesthesia by inserting it into the target bronchus through the nasal cavity or oral cavity via a bronchoscope. The fluid should be segmentally lavaged with 37 °C normal saline (1-2 ml/kg per session, with a total volume not exceeding 20 ml) and then recovered. The entire process must be strictly aseptic. When collecting blood, the appropriate site should be selected based on age (such as the anterior elbow vein of older children or the small veins of the hands and feet of infants). A 22-25G needle should be used for puncture (the tourniquet should not be used for more than one minute). After collecting, gently shake it to prevent blood clotting and mark it in time. The collection of cerebrospinal fluid requires a lumbar puncture[8]. The L3-L4 or L4-L5 intervertebral space is located in the lateral position of the child. After disinfection, the puncture needle is slowly inserted into the subarachnoid space. 1-3ml of cerebrospinal fluid is collected and aliquoting into sterile containers. During the operation, vital signs should be closely monitored, and the risk of brain herniation should be avoided. The BALF, Blood, and CSF were collected and maintained on ice for bacterial and fungal community analysis by amplicon-sequencing within 3 h. Basic information for patients is summarized in Table S1. The research purpose and consent were informed and duly signed by patients or their parents/legal guardians. Protocols and procedures in this study were approved by the Ethics Committee of Children’s Hospital Zhejiang University School of Medicine (Hangzhou), The Second Hosptial of Jiaxing (Jiaxing), Huzhou First People’s Hospital (Huzhou), Shaoxing Peoplès Hospital (Shaoxing), and Jinhua Maternal & Child Health Care Hospital (Jinhua).

### DNA extraction and nanopore sequencing

Samples were processed in the genetic diagnostic center. The BALF was collected and stored at 4°C before testing. The 7 reference strains and 13 clinically isolated strains were inoculated on Columbia blood plate culture medium for DNA extraction. These strains were identified by matrix assisted laser desorption ionization time-of-flight mass spectrometry (MALDI-TOF MS; Bruker Daltoniks, Germany) before DNA extraction. The QIAamp® DNA Mini Kit (QIAGEN, Germany) was used for the purification of microbial DNA. BALF (300mL) was first centrifuged at 13,000 rpm for 5 min to remove the supernatant, and then, the precipitate was suspended in normal saline (200 mL). The mixture was added to buffer ATL (180 mL) and proteinase K (20 mL), mixed by vortexing and incubated at 56 °C for 30 min at 500 rpm in a thermomixer (Eppendorf, Germany).

For the 21 bacterial strains, 200 mL of sample (1×108 CFU/ml), which was directly added to buffer ATL (180 mL) and proteinase K(20mL), was incubated at 56 °C for 30 min at 500 rpm in a thermomixer after mixing well. The following experimental steps were carried out according to the manufacturer’s instructions.

The concentration of isolated DNA from samples was measured with a Qubit dsDNA HS Assay Kit and a Qubit 4.0 Fluorometer (Thermo Fisher, USA) according to the instructions of the manufacturer. A total of 5∼10 ng of DNA template (10 mL) was used for 16S rRNA gene and ITS gene amplicon sequencing. The 27F/1492R and ITS1/4 primers were employed as the start primers for amplification of bacterial 16S rRNA and fungal internal transcribed spacer regions 1 and 2 (ITS1/2), respectively PCR amplicons were visualized by 1% agarose gel electrophoresis, followed by purification using 1x AMPure XP beads (Beckman Coulter, USA), and the DNA concentration was measured with a Qubit 4.0 Fluorometer. 1∼48 amplified DNA was pooled in equimolar amounts. The pooled DNA was end-repaired and dA-tailed with NEBNext FFPE DNA Repair Mix (New England Biolabs, USA) and NEBNext Ultra II End Repair/dA-Tailing Module (New England Biolabs, USA), followed by purification with AMPure XP beads (Beckman Coulter, USA). Then, the nanopore sequencing was performed using the ligation sequencing kit SQK-LSK110 (Oxford Nanopore Technologies, UK), and the DNA library was loaded into an R9.4.1 flow cell (Oxford Nanopore Technologies, UK) for sequencing for 1∼3 hours in the GridION platform according to the sample number. In every sequencing process run, negative process controls were added to the DNA extraction and amplification steps with sterile nuclease-free water to monitor contamination from the environment.

### Data preprocessing and amplicon analysis

All data were preprocessing and analysis on Baiyi Pathogenic Microorganism Bioinformatics Analysis Platform (Hangzhou Baiyi Technology Co., Ltd., Hangzhou, China). Briefly, the MinION data were transformed into FASTQ sequences with Guppy software. Subsequently, a comprehensive quality assessment and filtration process was carried out. Sequences that contained adapter and repetitive sequences, those with lengths either less than 200 bp or greater than 2000 bp, as well as sequences exhibiting a quality score Q lower than 7 were all removed. Microorganisms were classified as pathogens if they met the following criteria: having a relative abundance exceeding 0.01%, fulfilling a specific number of reads, and ranking within the Top 10. Specifically, the threshold for the number of specific reads was set at ≥ 3 reads for pathogenic bacteria and ≥ 1 read for fungi [9].

### Microbial community statistical analysis

Statistical analysis was conducted using R software. Measures were presented as mean ± standard deviation. The student’s t tests and ANOVA were employed for group comparisons. Cluster analysis was performed using the Ward’s Hierarchical Agglomerative Clustering Method (ward.D2) based on the Euclidean distance [10]. To identify differences in microbial characteristics by different characteristic factors, the Random Forest Model [11] was utilized, with a significance threshold value 0.05 and top 30 species. Genera with an abundance > 0.3% and present in more than 20% of the samples were selected for the construction of co-occurrences network based on the spearman’s correlation method [12]. The p value threshold for the correlation-based network was adjusted by ‘fdr’ method [13], and set as 0.05. And the correlation coefficient threshold for the correlation network was calculated using random matrix theory (RMT) based method [14]. The networks were visualized by using gephi [15]. Heatmaps illustrating the correlation between species and samples were generated using the pheatmap package (v 1.0.13) in R.

## Results

### Samples and characteristics of the data sets

In this study, we collected bronchoalveolar lavage fluid (BALF, n = 704), blood (n = 70) and cerebrospinal fluid (CSF, n = 8) samples from 782 children with severe infection in the early stage from the pediatric departments of 7 hospitals in the Hang-Jia-hu region, including 392 males and 364 females according to gender information. The original areas were Hangzhou (419 cases), Huzhou (54 cases), Jiaxing (34 cases), Jinhua (172 cases) and Shaoxing (50 cases). The symptoms were summarized as bronchitis (38 cases), bronchopneumonia (121 cases), cough (150 cases), fever (10 cases), pneumonia (194 cases) and sepsis (14 cases). According to the WHO age classification standard, the sample information was divided into Neonatal Period (< 1 month old, 3 cases), Infancy (< 1 year old, 67 cases), Toddlerhood (1-3 years old, 82 cases), and Preschool age (3-6 years old, 82 cases). 191 cases), School Age (6-12 years old, 231 cases), Adolescence (12-18 years old, 30 cases). BALF, blood and CSF samples were sequenced for 16S rRNA genes and ITS, respectively. In the end, we obtained better 16S rRNA gene sequencing results, but the ITS reads were too few to determine whether the corresponding fungal microorganisms were present in the sample, so in the end only a small part of ITS results is retained.

### Identification of the potential confounder in meta-analysis

In order to identify the influence of different classification of characteristic information on the composition and structure of microbial communities in samples, we analyzed the α diversity based on bacterial characteristic sequences according to the classification of characteristic information of cases. The abundance of bacterial species in BALF was significantly higher than that in blood and CSF (*p* < 0.05) (Figure 1-A). This suggests that the distribution of bacteria microbial community in BALF in the human body is affected by tissues and organs. Due to the large number of samples, the abundance of bacterial species in the samples from Hangzhou was significantly higher than that from Jiaxing, Huzhou, Shaoxing and Jinhua (*p* < 0.01) (Figure 1-C). At the same time, because the Jinhua area is geographically far away from Hangzhou, Huzhou, Jiaxing and Shaoxing, the bacterial diversity and evenness of the samples in Jinhua area were significantly different from those in Hangzhou-Jia-Hu area (*p* < 0.05) (Figure 1-D). The bacterial species in Toddlerhood samples were significantly different from those in Infancy, Adolescence, Neonatal Period, Preschool Age, and School Age samples (*p* < 0.05) (Figure 1-B). There were significant differences in diversity and evenness between Toddlerhood and School Age (*p* < 0.05). The bacterial species, diversity and evenness of Spesis samples were significantly lower than those of Bronchitis, Pneumonia and Bronchopneumonia samples (*p* < 0.05) (Figure 1-E, F). Moreover, there were no significant differences in microbial diversity and evenness or number of species among the three inflammatory samples. The number of bacterial species in the mildest cough samples was also significantly lower than that in the three types of inflammatory samples (*p* < 0.05).

**Figure 1.**
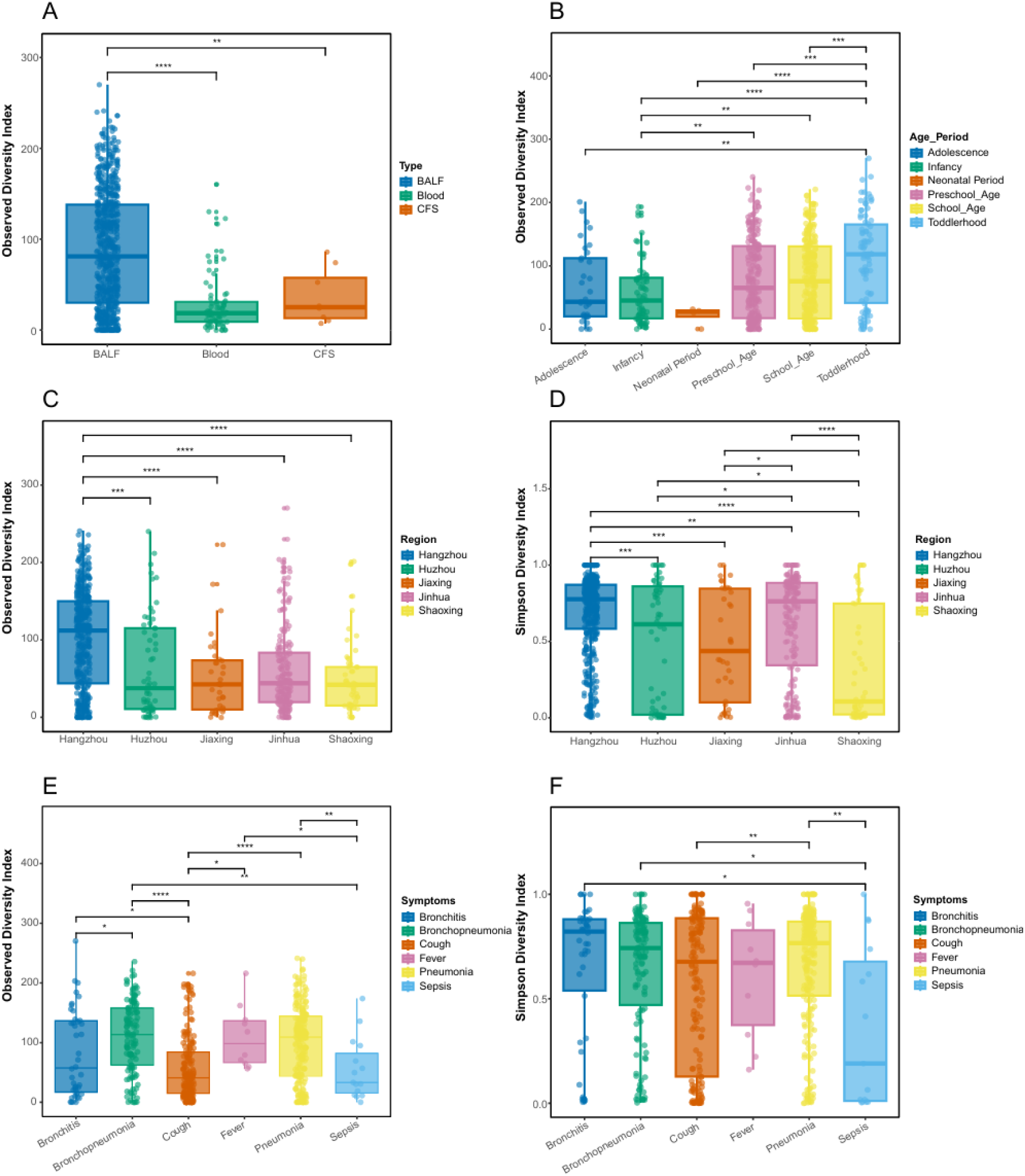
The box plots of the alpha diversity indexes of bacterial microbial community composition with different sample types, age periods, regions and symptoms. A and B shows the Observed diversity index of bacterial microbial communities in different sample types and regions separately. C and D represents the Observed and Simpson’s diversity index of bacterial microbial communities in different regions. E and F represent the Observed and Simpson’s diversity index of bacterial microbial communities with different symptoms. *, *p* < 0.05; **, *p* < 0.01; ***, *p* < 0.001.

Then, Dimension reduction Analysis based on Non-metric Multidimensional Scaling (NMDs) and community similarity test according to Analysis of Similarities using Permutation (Adonis) were performed on the microbial communities. The results showed that although NMDs dimension reduction analysis showed that the bacteria in BALF, CSF and Blood samples were not well separated (Figure 2-A), adonis test based on distance matrix showed that the bacterial community structure in BALF, CSF and Blood samples (Adonis, P < 0.05). p < 0.05) showed significant compositional differences (Table S4). Similarly, NMDs dimension reduction analysis showed that there was no significant difference in bacterial composition among the five regions (Figure 2-B). However, there were significant differences (Adonis, p < 0.05) in bacterial community composition between Hangzhou, jiaxing, huzhou, shaoxing and Jinhua. The microbial communities of hangzhou and the surrounding Jiaxing, Huzhou and Shaoxing were also significantly different (adonis, p < 0.05) (Table S3). In the BALF samples from Hangzhou area, the bacterial community composition in pneumonia cases and sepsis cases was significantly different from that in cough cases (adonis, p < 0.05) (Table S4). In the results of Adonis analysis, the explanatory variables of the model were less than 10%, indicating that the within-group variation in cough, pneumonia, and pyogenic cases was large, and there were other influencing factors of microbial composition. These results also have an assistant for explaining the low dispersion of NMDs (Figure S1-3).

**Figure 2.**
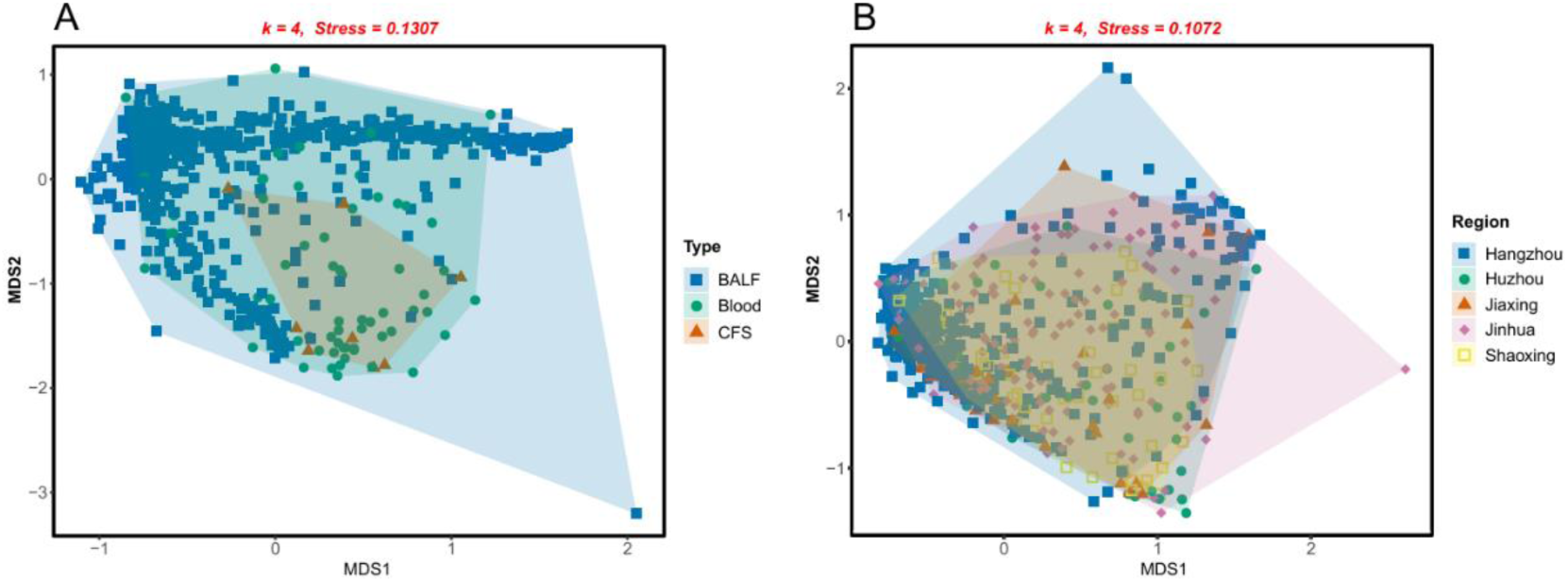
The dimensionality reduction analysis of bacterial microbial community composition with different sample types and regions. A shows the NMDs of bacterial microbial community composition in BALF, Blood, and CFS. B shows the NMDs of bacterial microbial community composition in different regions. The dimensionality number and ‘Stress’ parameters of the NMDs models were given and marked by red color.

### The cluster analysis of BALF hint characteristic bacteria in Hangzhou

We performed cluster analysis on BALF samples in Hangzhou and found that all samples could be divided into six clusters (Figure 3). Among them, CST-4 and CST-6 were enriched for *Chryseobacterium*, *Elizabethkingia*, *Flavobacterium*, *Epilithonimonas*, and *Kaistella* genera. In addition, CST4 enriched *Staphylococcus* and *Moraxella*, and it was distributed in all symptoms except cough samples, and its proportion was especially high in sepsis samples. It was the only cluster that exceeded CST-1 in the CST composition of each symptom. These results suggest that CST-4 may play an important role in the development of sepsis. CST-6 was also enriched for the genera *Weeksella* and *Paracidovorax* and was only found in pneumonia samples. The genera *Enterocloster*, *Hungatella*, and *Lacrimispora* were mainly enriched in CST-3, mainly in cough samples, and to a lesser degree in pneumonia and bronchopulmonary pneumonia, indicating that the proportion of the three enriched genera in CST-3 gradually decreased as respiratory symptoms worsened. *Bacteroides*, *Enterococcus*, *Neisseria*, *Veillonella*, *Streptococcus*, *Parabacteroides*, and *Porphyromonas* were enriched in the CST-2 cluster and in all symptom samples except sepsis samples. In the largest CST-1 cluster, *Staphylococcus*, *Streptococcus*, *Moraxella*, *Haemophilus*, *Comamonas*, and *Actinobacillus* genera were enriched and distributed in all symptoms. This suggests that the 12 genera enriched in CST-1 and CST-2 may play an important role in the development and progression of respiratory diseases.

**Figure 3.**
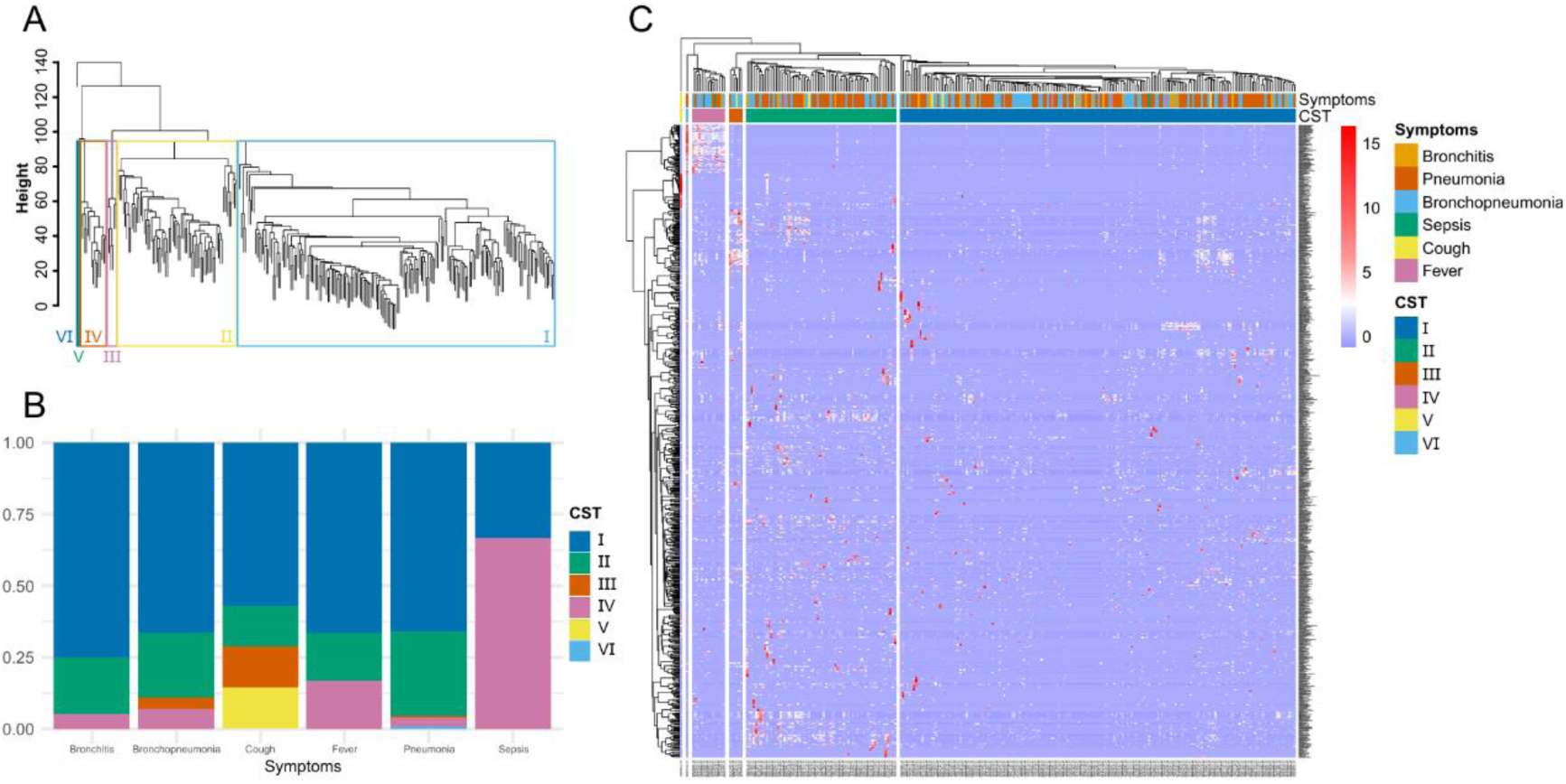
The clustering analysis for BALF in Hangzhou. A shows the six clusters based on Euclidean distance using the ward.D2 clustering method for the samples of BALF in Hangzhou. The different clusters were marked by colorful Roman numerals. B shows the distribution of the six clusters to which the samples of BALF belong in different symptoms. C shows the heatmap of species based on microbial abundance which was separated to six blocks based on Euclidean distance using the ward.D2 clustering method for the samples of BALF in Hangzhou.

**Figure 4.**
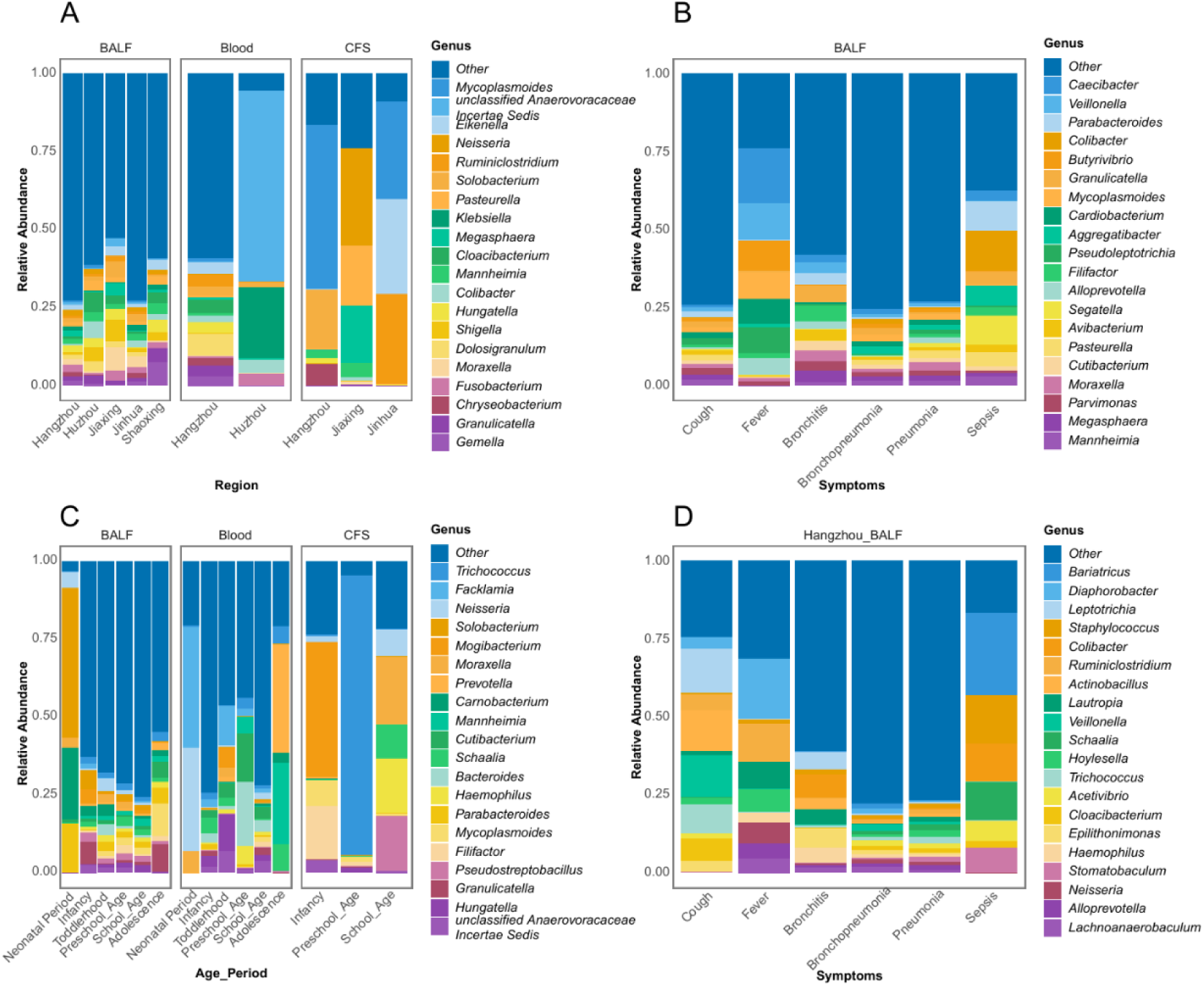
The stacked plots of microbial community composition at the genus level. A shows the microbial community composition at the genus level for different regions of different sample types. B shows the microbial community composition at the genus level for different symptoms of different sample types. C shows the microbial community composition at the genus level for different age periods of different sample types. D shows the microbial community composition at the genus level for different symptoms in the BALF of Hangzhou.

### Alterations of microbial composition in BALF samples

We analyzed the bacterial composition of all samples, and found that the sum relative abundance of the top 20 species in different regions was less than 50%, and the highest proportion of a single genus was only 7.5%, indicating that there was no dominant genus in the microbial community of BALF samples in different regions, and the community composition was uniform (Figure 3-A). The probiotic genus *Solobacterium*, opportunistic pathogen genus *Carnobacterium* and intestinal probiotic genus *Parabacteroides* were mainly detected in neonatal BALF (Figure 3-C). The results showed that the species of bacteria in the bronchoalveolar lavage fluid of neonatal children were relatively single, and the species of bacteria increased with age, and the distribution area was diversified. The sum of the relative abundance of the top 20 species in samples with fever and sepsis was 75% and 60%, respectively, while that in samples with other symptoms was less than 45%. The intestinal colonization genus *Caecibacter*, the respiratory colonization genus *Veillonella*, and the respiratory opportunistic pathogen genus *Mycoplasmoides* in fever samples, and the intestinal opportunistic pathogen genus *Colibacter* in pyogenic blood samples accounted for more than 10% (Figure 3-B). The sum of the relative abundance of the top 20 species with the highest proportion of bronchitis, bronchopulmonary pneumonia and pneumonia symptoms was less than 40%, and the sum of the relative abundance of the other three symptoms was more than 70%. Among them, the species composition of cough symptoms in BALF from Hangzhou was significantly different from the comprehensive situation in each region. The new genera with a relative abundance of more than 10% were *Leptotrichia* and *Actinobacillus* (Figure 3-D). unclassified *Dothideomycetes*, *Malassezia*, *Cladosporium*, *Catenaria*, and *Bjerkandera* accounted for a high proportion of the fungal composition in BALF samples from patients with cough and fever. In particular, the proportion of unclassified *Malasseziaceae* was as high as 70% in pneumonia cases (Figure S2). The opportunistic pathogen Ascotricha also accounted for more than 25% of the cases of bronchitis and bronchopneumonia.

### The identification of characteristic microbial community for BALF

We constructed robust random forest tree models to identify core communities and characteristic microorganisms in microbial communities. The results showed that although the characteristic organisms of pneumonia symptoms were not identified in BALF cases in Hangzhou area, However, the cough cases were characterized by oral pathogen *Actinomyces viscosus*, opportunistic pathogen *Capnocytophaga stomatis* and *Chroococcales cyanobacterium*. Fever cases were characterized by the opportunistic pathogens *Chitinophaga pinensis*, *Parafilimonas terrae* and *Aerococcus christensenii*. Sepsis was characterized by the opportunistic pathogen *Epilithonimonas hominis*, the intestinal probiotic *Coprobacter fastidiosus*, and the oral pathogen *Actinomyces oris* (Figure 5-A). Compared with cough and fever, the abundance of *Mycoplasma pneumoniae* and *Haemophilus parainfluenzae* in pneumonia and bronchopneumonia was significantly increased. *Haemophilus influenzae* was significantly more abundant in cases of pneumonia, bronchopneumonia, and bronchitis than in cases of sepsis, cough, and fever (Figure 5-B). The results showed that *Mycoplasma pneumoniae*, Haemophilus *parainfluenzae* and *Haemophilus influenzae* were the characteristic organisms of pneumonia, bronchopneumonia, bronchitis, cough, cold and pyogenic diseases in Hangzhou area. In addition, the random forest model could explain 49.13% of the differences in microbial communities caused by different regions, and identify the characteristic microorganisms associated with respiratory diseases in different regions. *Mycoplasmoides pneumoniae* was the characteristic microorganism of BALF microbial community in Shaoxing area, and the t-test showed that its abundance was significantly different from that in Jinhua and Hangzhou area (Figure S4-A). *Streptococcus*, *Haemophilus*, *Veillonella* and *Campylobacter* were the opportunistic pathogens that colonized the oral cavity, upper respiratory tract and gastrointestinal tract. And the t-test showed that the abundance of *Haemophilus parainfluenzae* was significantly higher than that of other areas (Figure S3-B).

**Figure 5.**
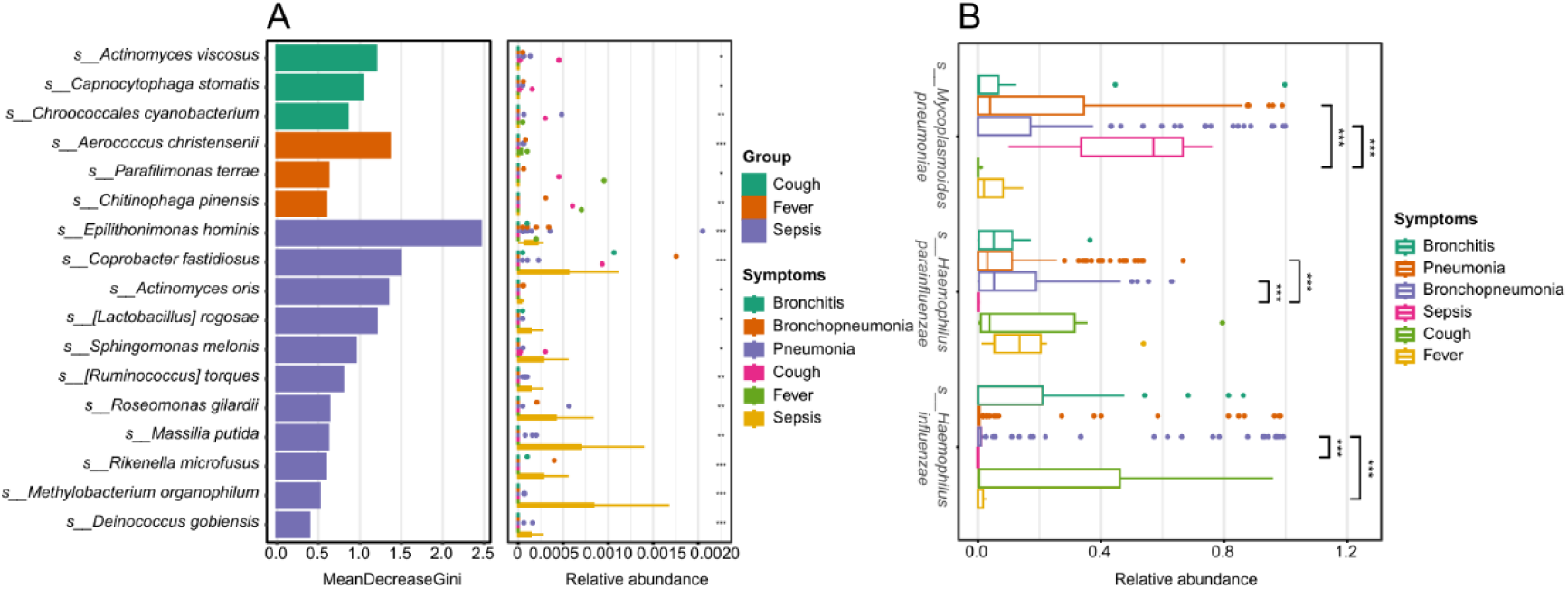
The statistical analysis charts of differential taxon by Random Forest analysis. A shows the Mean Decrease Gini index and relative abundance of characteristic species for BALF in Hangzhou filtered by the Random Forest model. B shows the differential species of symptoms for BALF in Hangzhou by Student’s T test. *, *p* < 0.05; **, *p* < 0.01; ***, *p* < 0.001.

### BALF, blood, and CSF have unique microbial co-occurrence network

The co-occurrence network was constructed based on the spearman correlation of microbial abundance, and the sub-networks were constructed according to different features of classification information. The topological parameters of each network were calculated to quantify the structure and characteristics of the network (Table-S5). The average degrees of all network nodes were higher than 10, which was higher than the average degrees of general microbial co-occurrence network [16]. Except for The average path length of The BALF network, which is greater than 6, The average path length of other sub-networks is less than 4. As we know, the smaller the average path length, the faster the material and information exchange between species. At the same time, the clustering coefficient reflecting the tightness of the network was also higher than 0.6, especially the clustering coefficient of the symptoms sub-network was as high as 0.81. These results suggest a very tight interaction between microorganisms in BALF samples. After the modularized calculation, the modularized coefficients are all over 0.8, indicating that all networks can be well modularized. In the sub-network (r > 0.54, p < 0.05) based on Symptoms cases in BALF samples in Hangzhou area (figure 6-A), a total of 51 node modules were obtained, among which the nodes of the first 15 modules accounted for 87.1% of all nodes. By tagging the pathogenicity of species nodes, we found that opportunistic pathogen nodes in module 1 (M1) and module 10 (M10) clustered into sub-modules and were closely connected with each other. This indicates that co-infection of these opportunistic pathogens may exist when conditions permit. We also found 11 module connectors (*Acinetobacter radioresistens*, *Desulfoscipio gibsoniae*, *Myxococcus fulvus*, Candidatus *Schmidhempelia bombi*, *Lelliottia amnigena*, *Staphylococcus carnosus*, *Herbaspirillum frisingense*, *Staphylococcus schleiferi*, *Peribacillus loiseleuriae*, *Staphylococcus arlettae*, *Apibacter mensalis*) and two module hubs (*Klebsiella michiganensis* and *Flavobacterium oncorhynchi*) (figure 6-A, B). Among them, opportunistic pathogens *Acinetobacter radioresistens* (M6, M10, and M11), *Staphylococcus schleiferi* (M1, M2, M5, and M7) and *Staphylococcus arlettae* (M1, M7 and M16) played as connectors for a total of 7 modules. *Klebsiella michiganensis*, as the core node in the module, is directly connected to most nodes in the module, including opportunistic pathogens, indicating that this species is highly correlated with the occurrence of other species in the module, and its existence plays an important role in the stability of the module network. In addition, we found that the three phyla of module connectors, *Pseudomonadota*, *Bacillota* and Bacteroidota, were also the three phyla with the largest number of node connections, accounting for about 85% of the total (Figure 6-C). These results indicated that the sub-network based on the Symptoms cases in BALF samples of severe children in Hangzhou could well show the co-occurrence relationship between microorganisms, especially pathogenic bacteria and opportunistic bacteria in BALF of severe children in Hangzhou area.

**Figure 6.**
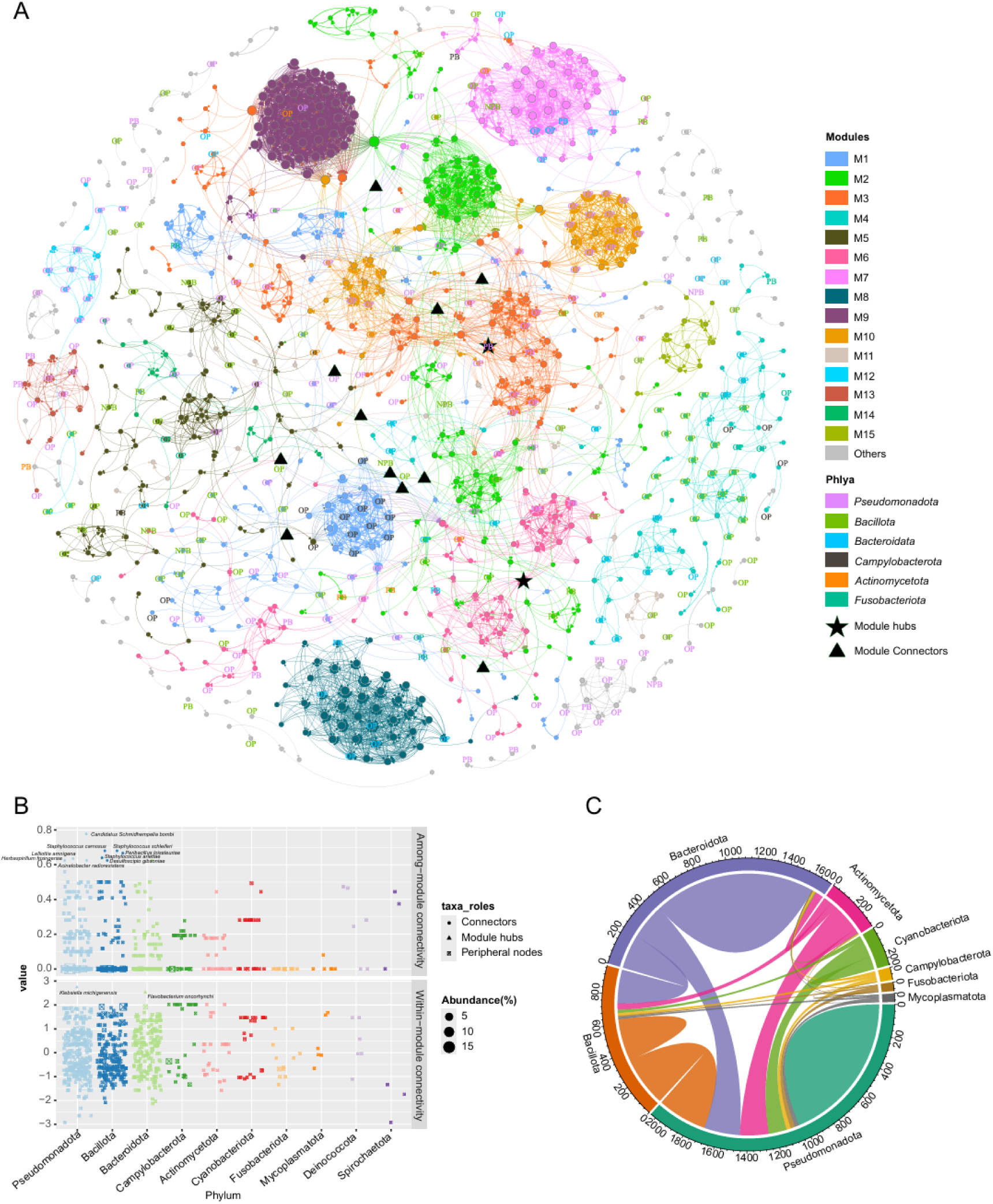
Microbial community ecological network and association analysis. A is the microbial ecological network, nodes are colored according to modules (M1-M15, etc.), node labels are colored according to Phyla classification, and shapes identify module roles (★ for module hubs, ▲ for module connectors). B is the scatter plot of module attribute-phylum association, showing the functional roles and abundance distribution of phyla classified microorganisms. C is the phylum level community node association chord diagram, which shows the number of species nodes among different phylum or within phylum by string connection.

## Discussion

In this study, we analyzed the microbial community amplicon-sequencing data from bronchoalveolar lavage fluid, blood, and cerebrospinal fluid of early stages of severe infection in children, explored microbial differences, and constructed a robust early stage of severe infection in children.

### The tissue specificity and pathological mechanism of microbial community distribution

We found that the abundance of bacterial species in BALF was significantly higher than that in blood and cerebrospinal fluid (p < 0.05), which was consistent with the physiological characteristics of the respiratory tract as an open system in direct contact with external pathogens. Respiratory mucosal barriers (such as mucus layer and ciliary motility) form a dynamic defense network with commensal microbiota. The diversity of bacterial flora in BALF may reflect the balance between microbial colonization and immune response during lower respiratory tract infection. For example, Staphylococcus, Moraxella and other genera were enriched in BALF. This is similar to a study of acute respiratory infections that found that children’s respiratory BALF was enriched for *Moraxella*, *Staphylococcus* and *Corynebacterium*, Nasal isolates of *Moraxella*, *Staphylococcus*, and *Corynebacterium* genus were also evaluated for their ability to induce epithelial damage and inflammatory responses [17]. The results showed that Moraxella catarrhalis induced significantly greater epithelial damage and inflammatory cytokine expression (IL-33 and IL-8), indicating a stronger ability to damage epithelial cells. These indicate that Staphylococcus and Moraxella may promote the progress of infection by destroying the tight junctions of epithelial cells [18]. The presence of Parabacteroides in neonatal BALF, A similar study evaluated the lung microbiome of bronchoalveolar lavage fluid (BALF) samples from children with hematological malignancies complicated with moderate to severe lower respiratory tract infections (LRTIs) and 21 LRTI children with matched age, sex, weight, and infection severity and no underlying malignancy [19]. The results showed that the α and β diversity of the lung microbiome in children with hematological malignancies and LRTIs were significantly reduced. The proportions of *Firmicutes*, *Bacteroidetes* and *Actinobacteria* significantly decreased. At the phylum level, *Proteobacteria* increased. The genus levels of *Paracbacteroides*, *Klebsiella*, *Gremonella*, *Escherichia*_*Shigella* were significantly higher than those of the control group. These results suggest that early respiratory microbiota may be regulated by gut-lung axis [20–22]. The high abundance of intestinal conditional pathogen *Colibacter* in blood samples (>10%) is associated with sepsis, suggesting that translocation of intestinal flora may be the underlying mechanism of severe infection [23]. The absence of ITS sequencing results in cerebrospinal fluid samples may be related to the low incidence of fungemia or insufficient sample size, but the risk of invasive fungi such as *cryptococcus* passing through the blood-brain barrier should be alert.

### The shaping effect of geographical and age factors on microbial communities

The differences in microbiota driven by geographical isolation shape the regional characteristics of epidemiology. The bacterial abundance of samples from Hangzhou was significantly higher than that of surrounding cities (p < 0.01), and the composition of bacterial flora in Jinhua area was significantly different from that in Hangzhou-Jia-hu area due to geographical distance (p < 0.05). This may be related to differences in regional air quality (e.g., PM2.5 concentration), distribution of healthcare resources (e.g., antibiotic use patterns), or exposure to environmental microorganisms [24]. Specific enrichment of the characteristic microorganism *Mycoplasmoides pneumoniae* in Shaoxing area. Similar studies found that the main transmission types of *Mycoplasmoides pneumoniae* in Japan in 2019-2020 were 2c and 2j, which had been spreading since 2010. There is local adaptation and selection pressure [25]. These results suggested that the local adaptive differences caused by the genotype differences of Mycoplasma pneumoniae strains in Shaoxing may also be related to the environmental selection pressure. The bacterial flora of neonatal BALF is dominated by probiotics such as Solobacterium, which gradually diversify with age, which is consistent with the maturation process of children’s immune system [26]. There was a significant difference in microbiota between Toddlerhood (1-3 years old) and school-age (6-12 years old) (p < 0.05), which may be related to exposure to the kindergarten/School environment, changes in dietary structure (such as complementary food addition), and vaccination history [27]. It is worth noting that the sample size of adolescents (12-18 years old) was small (30 cases), and whether their microbiota is similar to adult patterns still needs to be verified by larger samples.

### Symptom-specific microbiota characteristics are associated with disease progression

In this study, the proportion of Staphylococcus and Moraxella in CST-4 cluster was higher than that in CST-1 cluster. Considering that α -hemolysin heptamer of Staphylococcus can penetrate into the cell membrane of red blood cells, white blood cells and endothelial cells, and disrupt ion balance (such as Ca²⁺ influx), leading to osmotic pressure imbalance and lysis of cells. At the same time, α-hemolysin can be recognized by TLR2/TLR4 to release inflammatory factors. It recruited neutrophils and aggravated cell damage. They are phagocytosed by macrophages and release a large number of cytokines, forming a ’cytokine storm’ [28]. Thus, the high proportion of Staphylococcus and Moraxella species in spesis samples suggests that these bacteria may exacerbate systemic inflammatory responses by secreting toxins, such as Staphylococcus aureus α- hemolysin, or by activating TLR2/4 signaling. In addition, the diversity of bacterial flora in sepsis samples was significantly lower than that in inflammatory samples (p <0.05), which may reflect the immune escape phenomenon caused by dysbiosis of bacterial flora [29], that is, the excessive proliferation of dominant pathogenic bacteria inhibits other bacterial flora and forms a "niche monopoly". With the progress of respiratory symptoms, the microbiota also showed a succession rule from oral colonization to respiratory pathogens. *Leptotrichia* (oral opportunistic pathogen) and Actinobacillus were enriched in cough samples, while Mycoplasma pneumoniae and *Haemophilus parainfluenzae* were increased in pneumonia samples. It is suggested that the evolution of microbiota from mild respiratory infection to severe pneumonia may follow the pattern of "oral bacterial colonization → respiratory pathogenic bacteria dominance"[30, 31]. However, there is no significant difference in the microbiota between bronchitis, bronchopulmonary pneumonia and pneumonia, which may indicate that these three diseases have overlapping pathological mechanisms at the microbial level [32], and need to be further distinguished in combination with host immune indicators. This result is consistent with the results of Wu and Segal (2018) et al on the mechanism of microbial community imbalance in the lower respiratory tract. This study suggested that the microbial communities of bronchitis, bronchoponia and pneumonia were characterized by growth and decreased diversity of potential pathogenic bacteria [33].

### Microbial co-occurrence networks and synergistic mechanisms of infection

The topological characteristics and potential functions of the co-infection network of opportunistic pathogens were identified by constructing the community co-occurrence network. The opportunistic pathogen nodes of module 1 (M1) and M10 in the co-occurrence network were closely connected, such as *Acinetobacter radioresistens* and *Staphylococcus schleiferi* as module connectors. Virulence gene expression may be coregulated by secreted quorum-sensing molecules, such as AHLs [34–37]. *Klebsiella michiganensis*, the core node, is directly connected to several opportunistic pathogens, and the resistant plasmids carried by it, such as blaKPC, may accelerate the spread of multidrug-resistant bacteria through horizontal transfer [38–40]. Based on the microbiota interaction network, it is possible to provide regulatory-based intervention in the early stage of severe pediatric disease from a clinical perspective. 85% of the connections in the network were concentrated in the three phyla *Pseudomonadota*, *Bacillota*, and *Bacteroidota*, suggesting that ecological regulation targeting these dominant phyla may be more likely to reshape the balance of the microbiota. For example, specific clearance of *Staphylococcus* using phage therapy [41, 42], or competitive inhibition of *Pathobateroides* colonization by probiotics such as Bifidobacterium [43–45], may be an adjuvant treatment strategy for severe infections.

## Conclusion

The respiratory tract related diseases are the main causes of death in children. Child pulmonary severe infection is the most important inducement for Respiratory tract related diseases. Therefore, it is necessary to understand the composition of the microbial community and the species correlation in children with severe pulmonary infection. In this study, the bronchoalveolar lavage fluid (BALF), blood and cerebrospinal fluid (CSF) samples of 782 children with severe infection in the early stage were systematically analyzed by amplicon-sequencing technology to reveal the distribution of microbial community and its clinical correlation. The results showed that the abundance of bacterial species in bronchoalveolar lavage fluid was significantly higher than that in blood and cerebrospinal fluid, and the community structure was regulated by the microenvironment of tissues and organs. Geographical differences led to significant differences in the composition of microbiota between Hangzhou and Jinhua. Shaoxing had enrichment of characteristic microorganisms such as Mycoplasma pneumoniae. Age affected the diversity of microbiota. In neonatal samples, probiotics such as *Solobacterium* were the main bacteria. The structure of microbiota gradually became complex with age, and there was a significant difference between children aged 1-3 years and 6-12 years. The diversity of microbiota in sepsis samples was lower than that in inflammatory diseases, and Staphylococcus and Moraxella were prominent in CST-4 cluster. The progression of respiratory symptoms is accompanied by the bacterial succession from oral colonizers to respiratory pathogens. The microbial co-occurrence network analyzed by third-generation sequencing technology showed the synergistic effect of opportunistic pathogens, which provided a theoretical basis for ecological targeted therapy.

